## Supplementary Information for "Robust proteome profiling of cysteine-reactive fragments using label-free chemoproteomics"

### Supplementary Figures

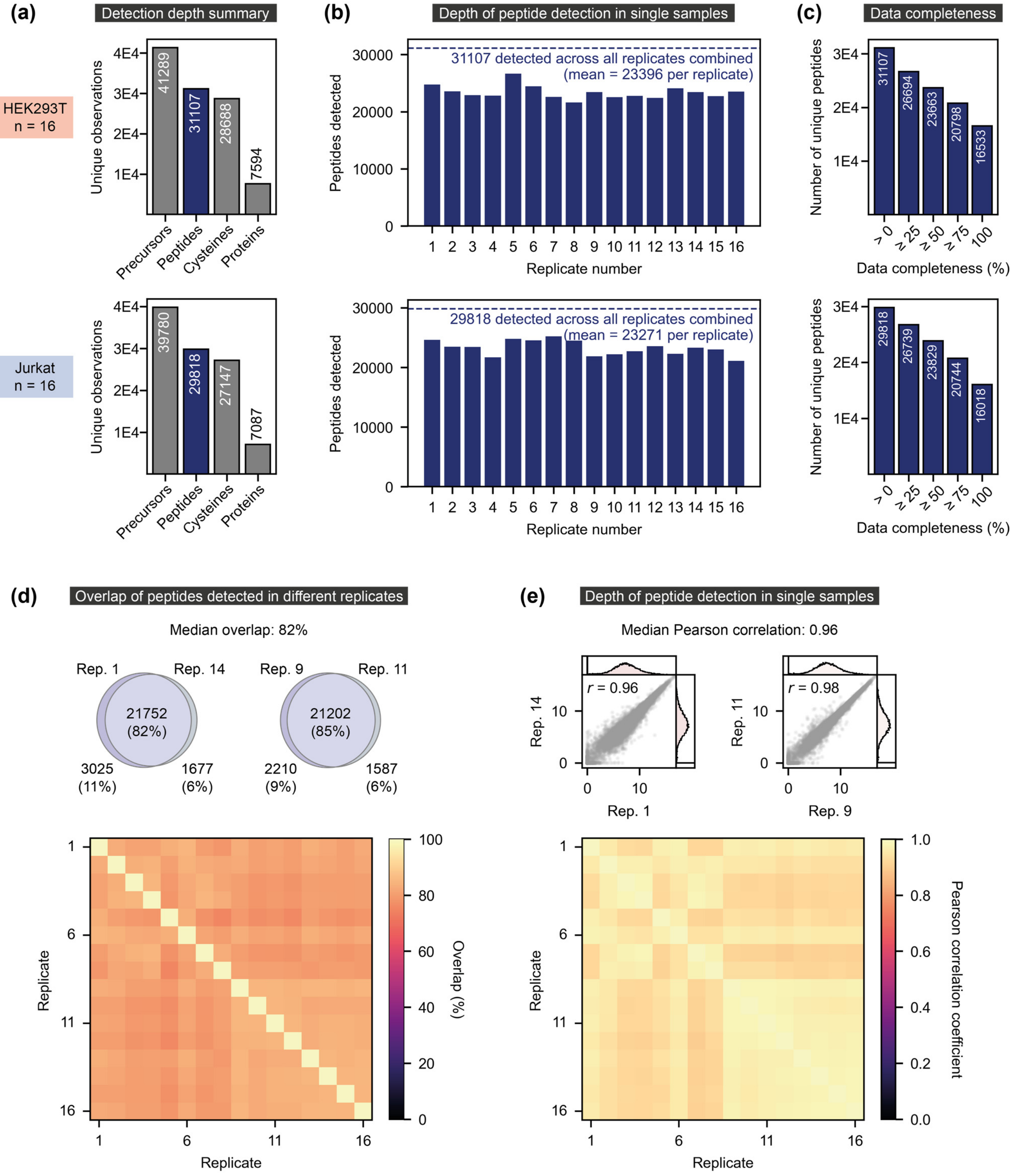

**Supplementary Figure 1. Data depth and completeness.** The detection depth and completeness of chemoproteomics data from HEK293T and Jurkat cell lysate. Data was obtained using 16 replicate DMSO control samples prepared from each lysate, and is summarised based on the total detection depth across all replicates **(a)**, the number of peptides detected in each individual replicate sample **(b)**, and the number of peptides detected at varying levels of data completeness **(c)**. In **(d)**-**(e)**, a more detailed view of peptide detection reproducibility is shown for the HEK293T lysate data, showing both the overlap in the peptides detected by each replicate (median overlap of 82% between any two replicates) and the correlation between the ($log_2$-transformed) intensity of these peptides (median Pearson correlation between any two replicates of 0.96). Heatmaps summarise the pairwise similarity between replicates whilst Venn diagrams of detected peptides **(d)** and correlation plots of $log_2$-transformed peptide intensities **(e)** are shown for representative replicates at the top of both figures.

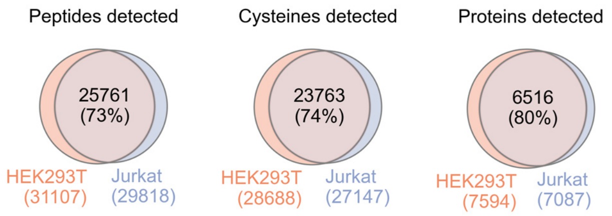

**Supplementary Figure 2. Overlap of peptide, cysteine, and protein detection in HEK293T and Jurkat lysates.** Data was obtained using 16 replicate DMSO control samples from each lysate.

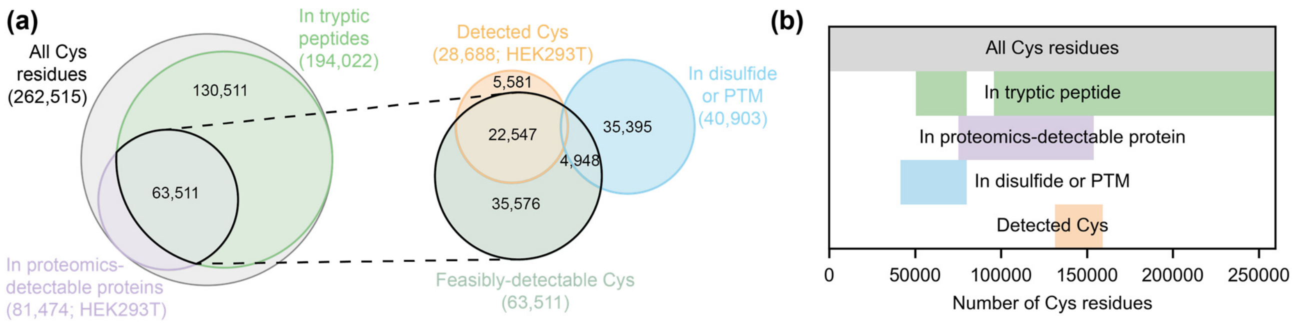

**Supplementary Figure 3. Estimating the size of the cysteinome that is 'feasibly-detectable' using IA-DTB chemoproteomics.** Data is summarised both through Venn diagrams **(a)** and a supervenn plot **(b)**.[1] Cysteine residues that are detected through IA-DTB chemoproteomics must have fulfilled two criteria: they were present within tryptic peptides of a suitable size for detection by mass spectrometry and they were present in proteins that are sufficiently abundant in the cell lysate. We have performed an *in silico* trypsin digestion of all proteins (~20,000) in the human proteome and estimate that, of the ~263,000 unique cysteine residues (grey), ~194,000 of these lie within tryptic peptides of an appropriate size (see experimental methods) (green). We have approximated the number of cysteines present in proteins of sufficient abundance in HEK293T lysate by assessing which proteins can be detected by standard proteomics methods, which accounted for ~81,000 cysteines (purple). Taken together, this gives an estimate of ~64,000 cysteine residues that lie within tryptic sequences of sufficiently abundant proteins in HEK293T lysate. Excluding the ~40,000 cysteine residues that are annotated in UniProt as being involved in disulfide bonds or other post-translational modifications (blue), our IA-DTB chemoproteomics experiments performed in HEK293T lysate (n=16 DMSO control samples) detect ~40% of the 'feasibly-detectable' cysteine residues (orange).

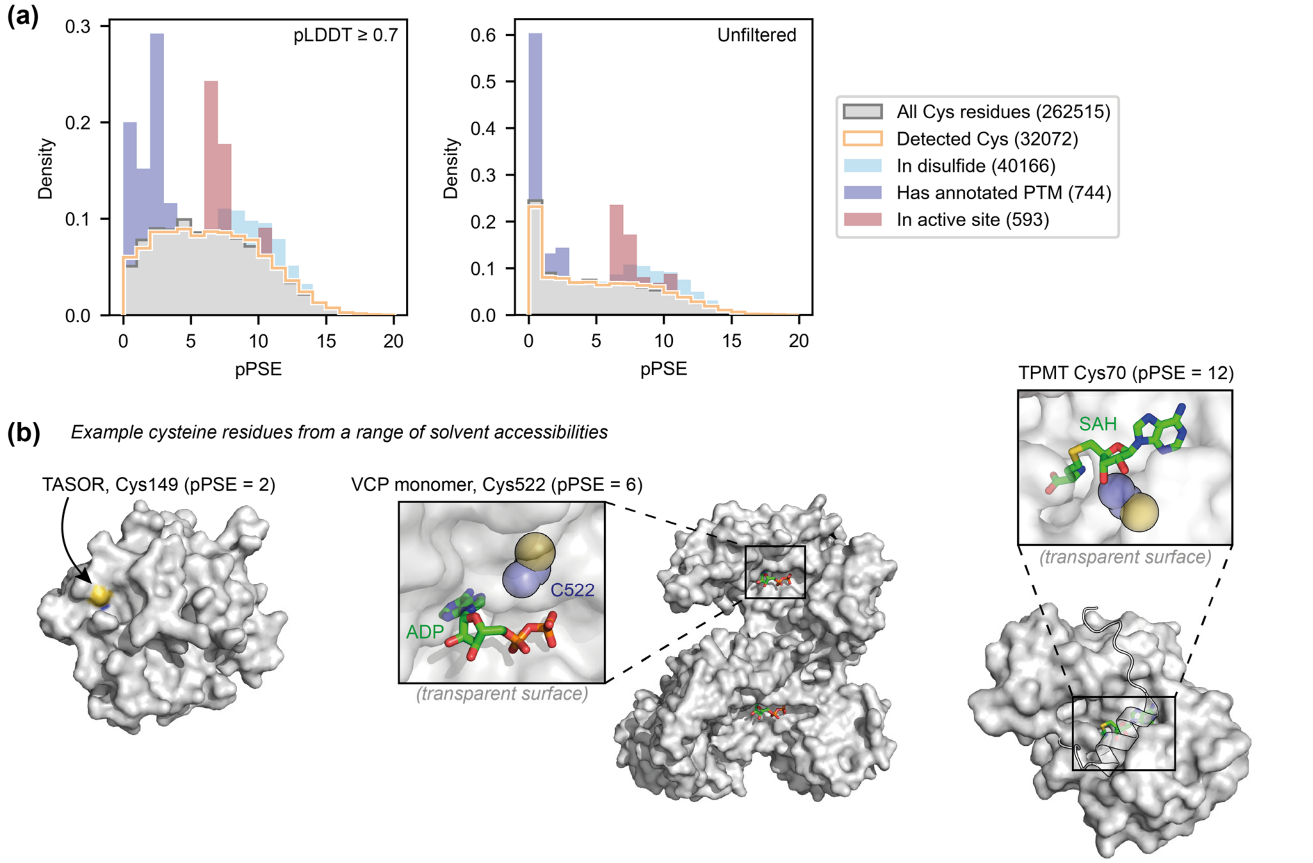

**Supplementary Figure 4. Surface exposure of residues detected through the IA-DTB probe in HEK293T lysate.** We detect cysteine residues from a range of solvent accessibilities (quantified using the pPSE metric) (as shown in the histogram in **(a)**),[2] including fully solvent-exposed residues (*e.g.*, TASOR Cys149, pPSE 2; PDB 6TL1),[3] residues in metabolite binding pockets (*e.g.*, VCP Cys522, pPSE 6; PDB 5FTJ),[4] and residues that lie deep within binding channels (*e.g.*, TPMT Cys70, pPSE 12; PDB 2H11; cartoon representation used for residues 17-39) **(b)**.[5] While pPSE values were calculated using AlphaFold2 models, the protein structures shown are from experimentally validated structures.[6]

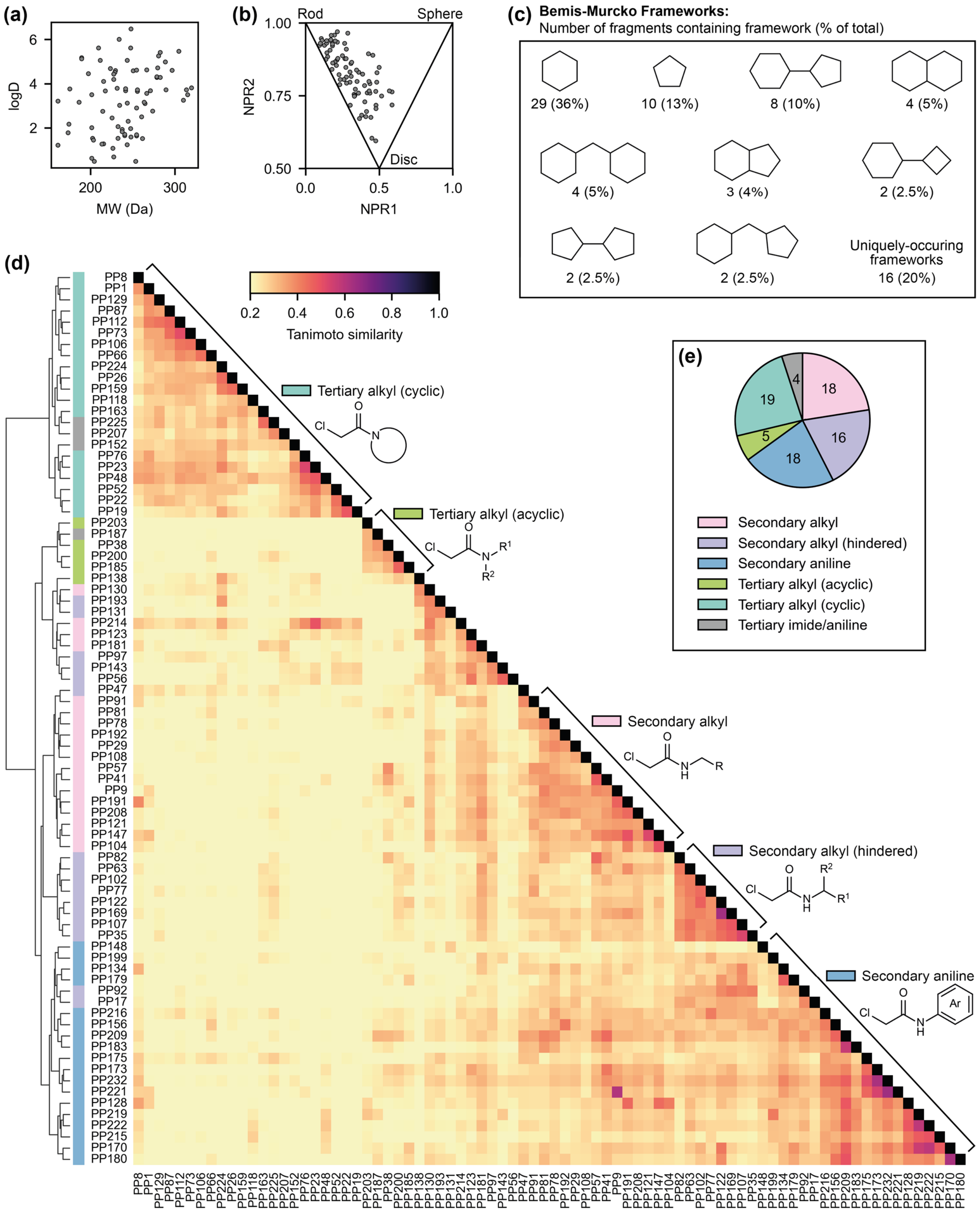

**Supplementary Figure 5. Chemical and physical properties of our 80-member library of chloroacetamide-functionalised fragments. (a)** All compounds possess a molecular weight that is consistent with the size range typically recommended for small molecule fragments.[7] **(b)** The library as a whole shows a diversity of physical shapes, based on the normalised principal moment of inertia ratios (NPR) obtained for these fragments from their predicted three-dimensional structures.[8] **(c)** As the size limit of small molecule fragments places restraints on the types of molecular frameworks that can be explored, the vast majority of compounds in our library consist of a small number of frameworks. However, the vast majority of the scaffolds in our library have previously been identified as structures that are well-suited to binding proteins.[9] **(d)** Tanimoto similarity matrix, based on Morgan molecular fingerprints, with hierarchical clustering. In general, there is little similarity between library members, and the clusters of similarity that are detected are largely driven by structural features adjacent to the chloroacetamide functional group. **(e)** Pie chart illustrating the distribution of structural features adjacent to the chloroacetamide functional group.

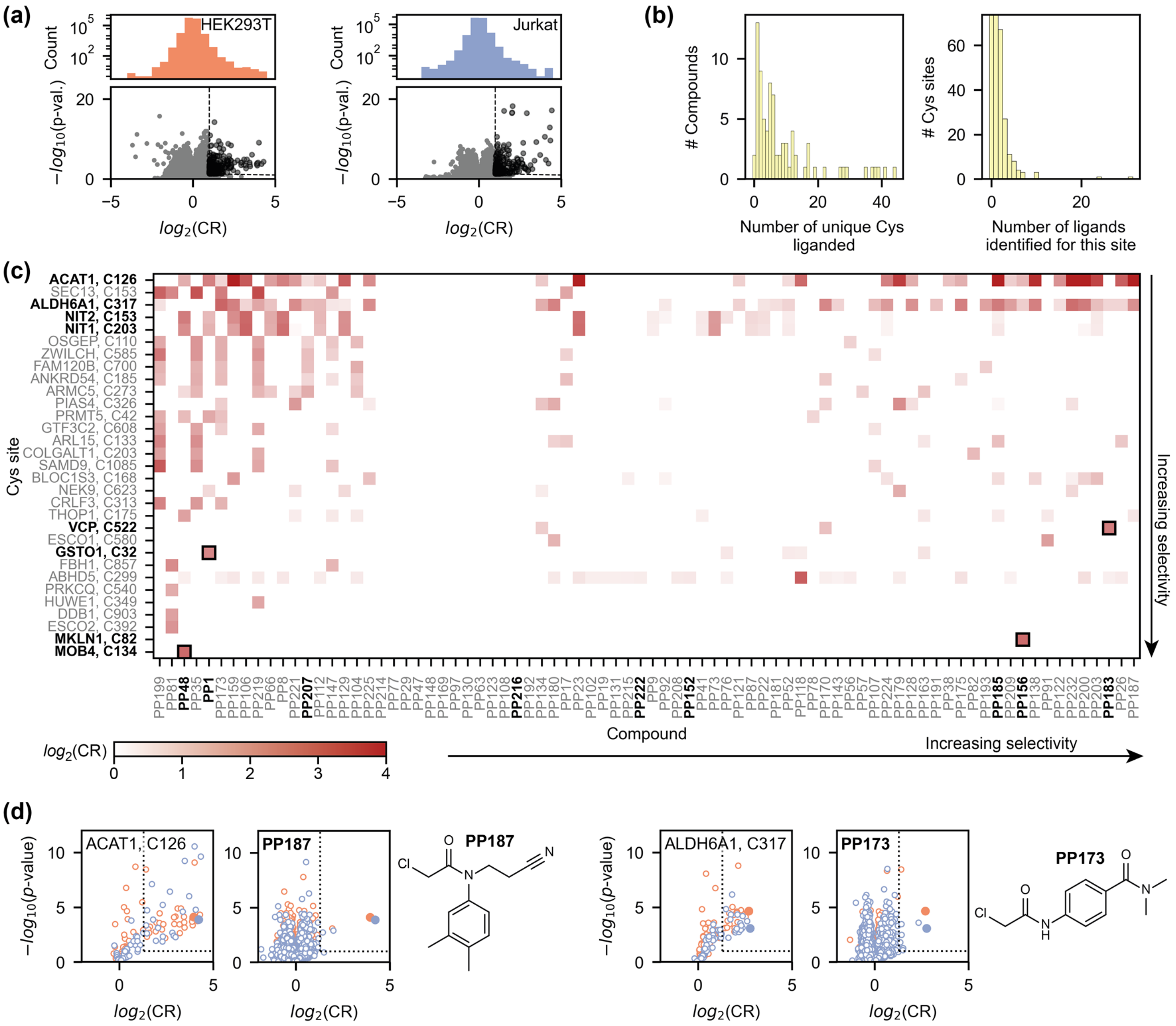

**Supplementary Figure 6. Data from the 80-compound screen performed in HEK293T and Jurkat lysate.** **(a)** Volcano plot and *log*2(CR) distributions of all the data obtained in HEK293T (left) and Jurkat (right) lysate (n=4 for compound-treated samples, n=16 for DMSO control samples). Each data point reports on the binding of an individual compound (50 µM) to an individual cysteine-containing peptide. **(b)** Distribution of compound (left) and cysteine (right) promiscuity. **(c)** Heatmap of all the interactions detected in Jurkat lysate. For clarity, this heatmap only includes cysteines that are liganded by at least compound with *log*2(CR) ≥ 1.5. Selective interactions are highlighted by black boxes, and the compounds/Cys sites that are discussed in the main text are labelled in bold. **(d)** Volcano plots for two active site cysteine residues (ACAT1 Cys126 and ALDH6A1 Cys317). While these cysteine residues bind many compounds, two compounds were identified that only bind these cysteine sites (PP187 and PP173, respectively). HEK293T data: orange; Jurkat data: blue.

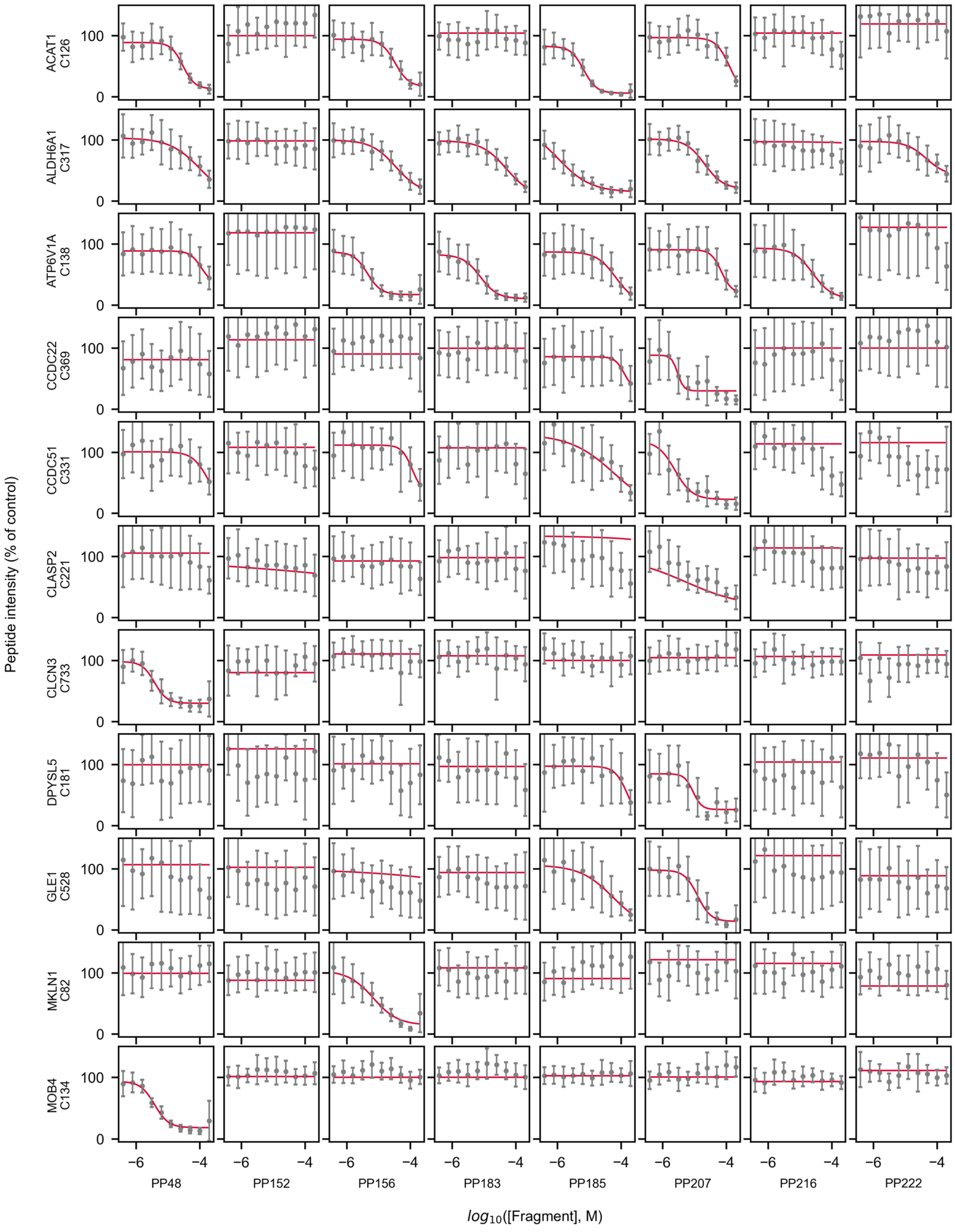

**Supplementary Figure 7. Concentration-response curves for all peptides with concentration-dependent interactions.** Continued over the page.

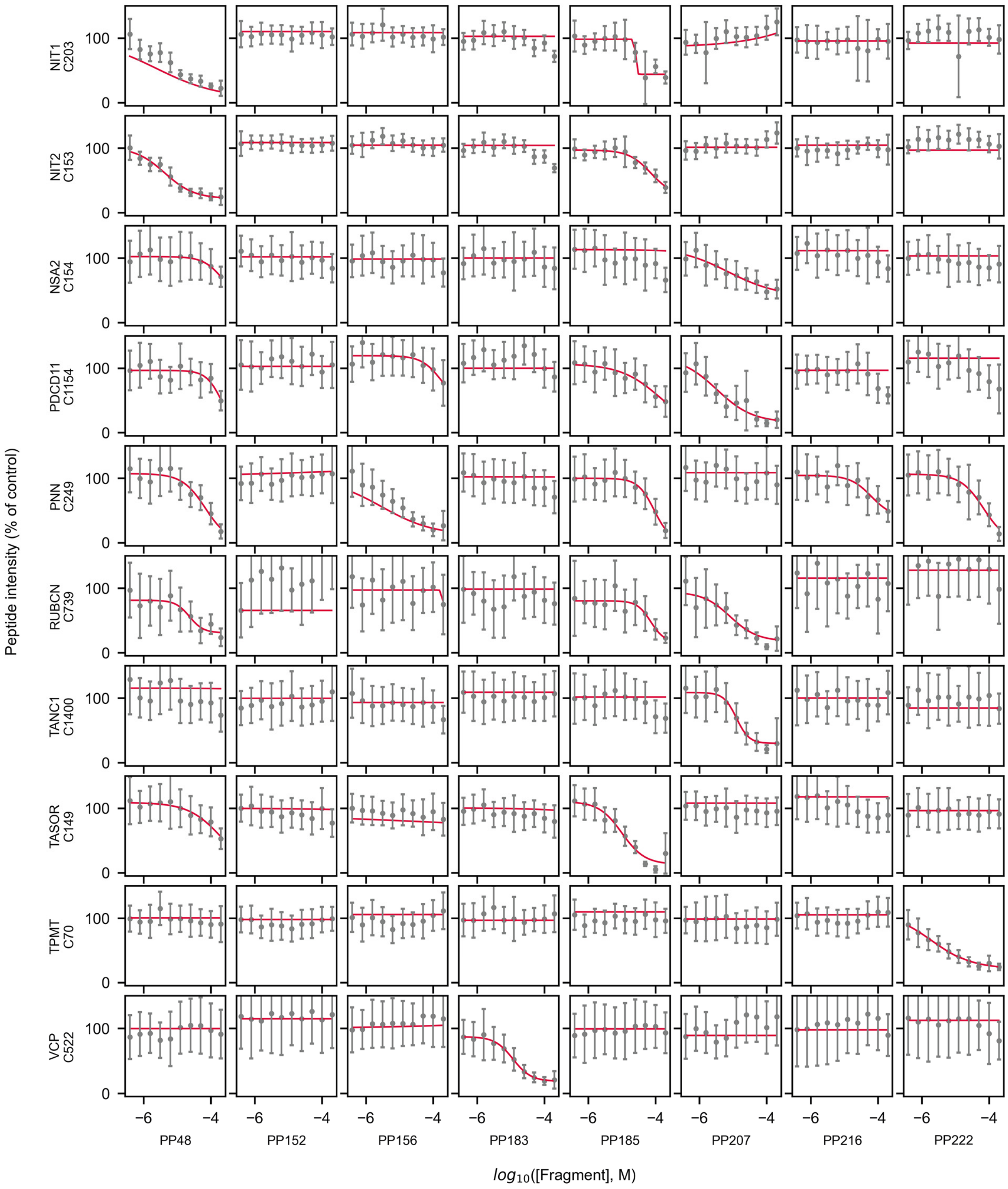

**Supplementary Figure 7. Concentration-response curves for all peptides with concentration-dependent interactions.** Data points represent the mean peptide intensity (relative to the DMSO control; error bars represent the standard deviation) measured at varying concentration of eight different chloroacetamides. Red lines represent the results of fitting a four-parameter logistic function to the data, which was used to identify concentration-dependent interactions. The resulting $pTE_{50}$ values are summarised by the heatmap in **Fig. 4e**. Compound treatments were performed in HEK293T lysate (n=4 for compound-treated samples, n=25 for DMSO control samples).

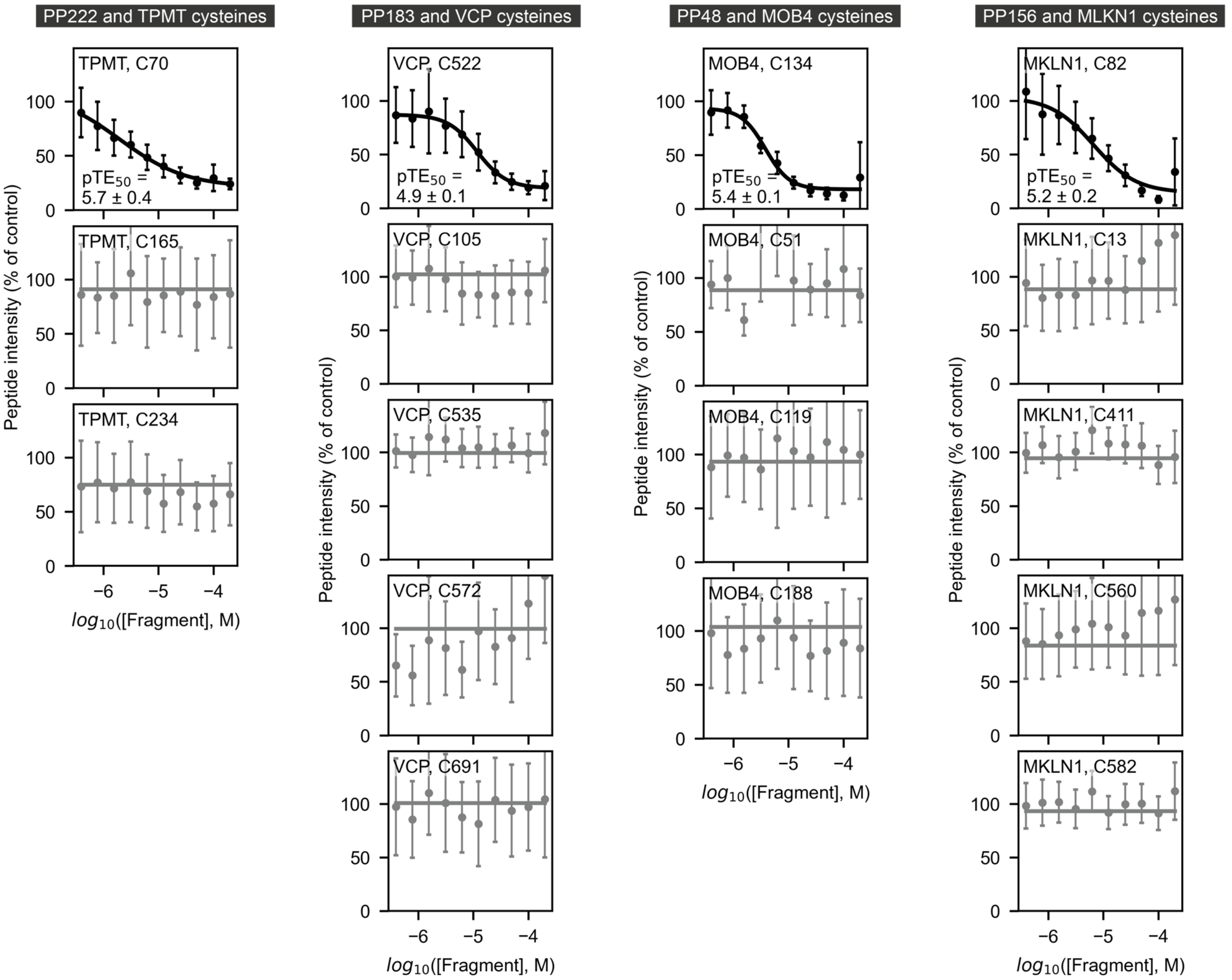

**Supplementary Figure 8. Concentration-response curves for all detected cysteine residues in TPMT, VCP, MOB4, and MKLN1 with the hit fragments for these proteins.** Data points and error bars represent the mean peptide intensity and standard deviation, respectively. Compound treatments were performed in HEK293T lysate (n=4 for compound-treated samples, n=25 for DMSO control samples).

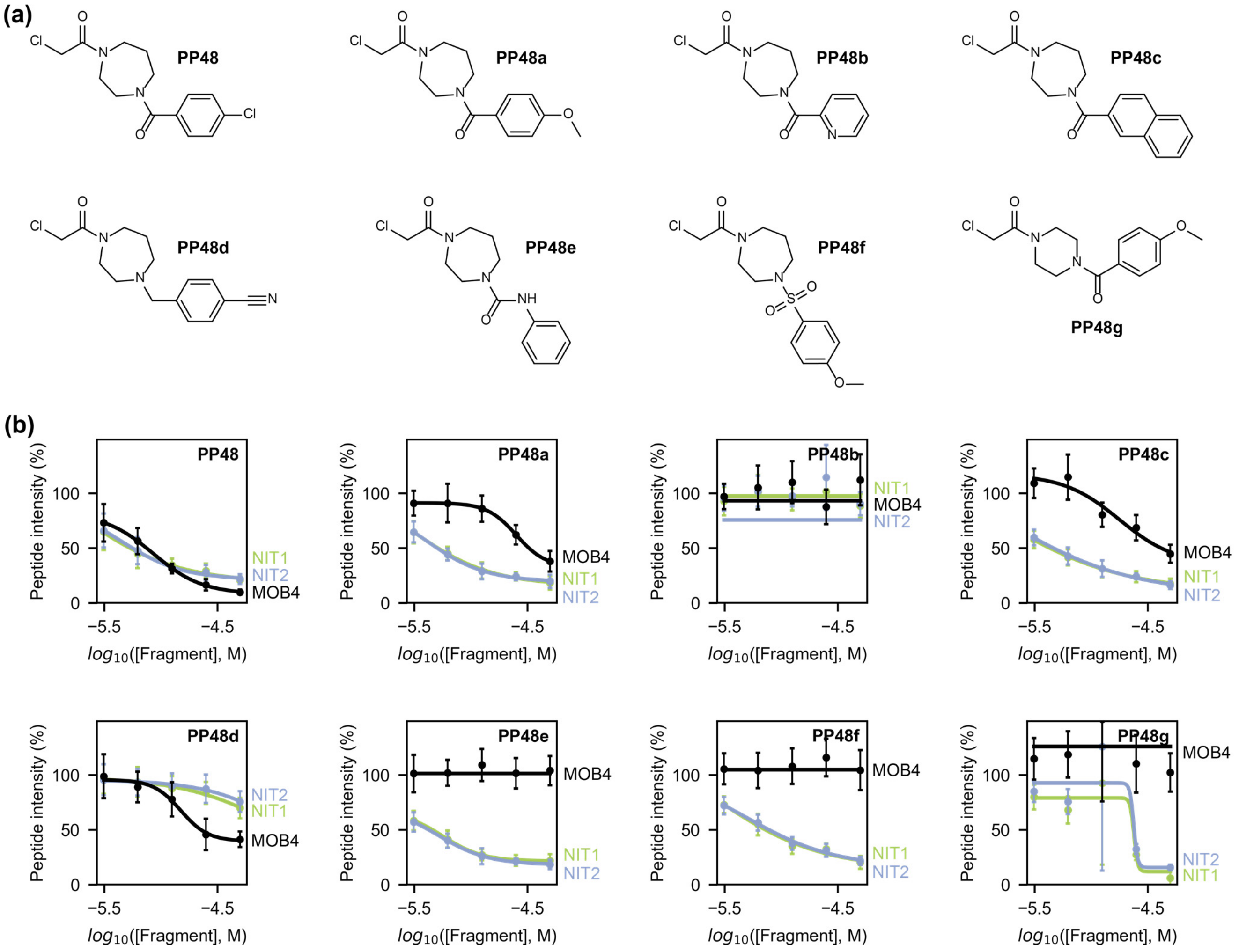

| Cys target | PP48 | PP48a | PP48b | PP48c | PP48d | PP48e | PP48f | PP48g |
|---|---|---|---|---|---|---|---|---|
| MOB4 Cys134 | 5.1 ± 0.1 | 4.6 ± 0.1 | NB | 4.8 ± 0.1 | 4.8 ± 0.1 | NB | NB | NB |
| NIT1 Cys203 | 5.7 ± NE | 5.5 ± 0.9 | NB | 5.8 ± NE | 3.9 ± NE | 5.2 ± 0.1 | 5.6 ± NE | 4.6 ± NE |
| NIT2 Cys153 | 5.4 ± 0.6 | 5.4 ± 0.4 | NB | 5.8 ± NE | 3.8 ± NE | 5.3 ± 0.2 | 5.3 ± 0.4 | 4.6 ± NE |

**Supplementary Figure 9. MOB4, NIT1, and NIT2 concentration-response data for a panel of compounds derived from PP48. (a)** Structures of the seven analogues of PP48 that were used to explore SAR around binding to MOB4, NIT1, and NIT2. **(b)** Concentration-response data are shown as the mean ± standard deviation for each compound concentration alongside the logistic regression curve. The $pTE_{50}$ values and associated errors from this regression analysis are shown in the table. Compound treatments were performed in HEK293T lysate (n=4 for compound-treated samples, n=32 for DMSO control samples). NB: no binding detected. NE ('not estimated') refers to errors whose value could not be accurately estimated.

### Supplementary Tables

**Supplementary Table 1. SMILES for all chloroacetamides used in this study.**

| Compound number | SMILES |
|---|---|
| PP1 | Fc1ccc(cc1)S(=O)(=O)C1CCN(CC1)C(=O)CCl |
| PP8 | Fc1cccc(NC(=O)CN2CCN(CC2)C(=O)CCl)c1 |
| PP9 | CS(=O)(=O)c1ccc(CNC(=O)CCl)cc1 |
| PP17 | CC(Cc1cc(C)[nH]n1)NC(=O)CCl |
| PP19 | ClCC(=O)N1CCCCC11CCOCC1 |
| PP22 | ClCC(=O)N1CC2(C1)CCOCC2 |
| PP23 | ClCC(=O)N1CCN(CC1)C(=O)N1CCCCC1 |
| PP26 | ClCC(=O)N1CCCCCC1c1ncon1 |
| PP29 | CCc1c[nH]c(CNC(=O)CCl)n1 |
| PP35 | CC(NC(=O)CCl)c1noc(Cc2ccccc2)n1 |
| PP38 | Cc1ccc(CN(CCO)C(=O)CCl)cc1 |
| PP41 | Cc1ccc(cc1)C(=O)NCCNC(=O)CCl |
| PP47 | Cc1cccc(c1)N1CCC(NC(=O)CCl)C1=O |
| PP48 | ClCC(=O)N1CCCN(CC1)C(=O)c1ccc(Cl)cc1 |
| PP52 | ClCC(=O)N1CCOC2(CCC2)CC1 |
| PP56 | CNC(=O)C1(CCCCC1)NC(=O)CCl |
| PP57 | Cc1ccc(CC(CO)CNC(=O)CCl)cc1 |
| PP63 | NC(=O)C(Cc1ccccc1)NC(=O)CCl |
| PP66 | Cc1conc1C1CN(C1)C(=O)CCl |
| PP73 | ClCC(=O)N1CCCC(C1)c1ncc[nH]1 |
| PP76 | CS(=O)(=O)N1CCCN(CC1)C(=O)CCl |
| PP77 | Cn1nccc1C(NC(=O)CCl)c1ccccc1 |
| PP78 | Cc1ccn2cc(CNC(=O)CCl)nc2c1 |
| PP81 | Cc1ccc(C)c(Oc2ccc(CNC(=O)CCl)cn2)c1 |
| PP82 | Cc1ccc(CC(NC(=O)CCl)c2ccccn2)cc1 |
| PP87 | ClCC(=O)N1CCC(Cc2nccs2)C1 |
| PP91 | Cc1cccc(c1)C(CNC(=O)CCl)N1CCOCC1 |
| PP92 | CC(NC(=O)CCl)c1ccc2[nH]ncc2c1 |
| PP97 | ClCC(=O)NC1(CCC1)c1ncccn1 |
| PP102 | COCC[C@@H](NC(=O)CCl)c1ccccc1 |
| PP104 | Fc1cnc(CCNC(=O)CCl)c(F)c1 |
| PP106 | ClCC(=O)N1CC(C1)c1ccc(Br)cn1 |
| PP107 | CC(NC(=O)CCl)c1csc(n1)-c1ccccc1 |
| PP108 | Oc1nc2CCCCc2cc1CNC(=O)CCl |
| PP112 | ClCC(=O)N1CCC(C1)c1ccn[nH]1 |
| PP118 | ClCC(=O)N1CCc2nc[nH]c2C1c1ccc(Cl)cc1 |
| PP121 | ClCC(=O)NCCc1ccc2OCCOc2c1 |
| PP122 | COc1ccccc1C(C)NC(=O)CCl |
| PP123 | CN(C)C1(CNC(=O)CCl)CCCCC1 |
| PP128 | Fc1ccc(NC(=O)CCl)c(F)c1 |
| PP129 | ClCC(=O)N1CCC(CC1)c1nc2ccccc2o1 |
| PP130 | ClCC(=O)NCC1CCC1 |
| PP131 | FC(F)(F)C1CCC(CC1)NC(=O)CCl |
| PP134 | Cc1nc(cs1)-c1cccc(NC(=O)CCl)c1 |

| | |
|---|---|
| PP138 | CN(C1CCCCC1)C(=O)CCl |
| PP143 | ClCC(=O)NC1(CCCC1)C#N |
| PP147 | Fc1ccc(CCNC(=O)CCl)cc1 |
| PP148 | FC(F)(F)Cc1nnc(NC(=O)CCl)o1 |
| PP152 | ClCC(=O)N1CCCC1=O |
| PP156 | Cc1cc(NC(=O)CCl)no1 |
| PP159 | ClCC(=O)N1CCCC1c1ccsc1 |
| PP163 | OC(=O)C1Cc2ccccc2CN1C(=O)CCl |
| PP169 | CC(NC(=O)CCl)c1cccc2ccccc12 |
| PP170 | COc1ccc(Cl)cc1NC(=O)CCl |
| PP173 | CN(C)C(=O)c1ccc(NC(=O)CCl)cc1 |
| PP175 | CC1CN(CC(C)O1)c1ccc(NC(=O)CCl)cc1 |
| PP179 | COc1ccc(cc1)-c1csc(NC(=O)CCl)n1 |
| PP180 | COc1cc(Cl)c(C)cc1NC(=O)CCl |
| PP181 | ClCC(=O)NCC1(CC1)c1ccccc1 |
| PP183 | Cc1nnc([nH]1)-c1ccc(C)cc1NC(=O)CCl |
| PP185 | COc1cccc(CN(C)C(=O)CCl)c1 |
| PP187 | Cc1ccc(cc1C)N(CCC#N)C(=O)CCl |
| PP191 | Fc1cccc(CNC(=O)CCl)c1 |
| PP192 | Cc1noc(C)c1CNC(=O)CCl |
| PP193 | CC1CCCCC1NC(=O)CCl |
| PP199 | ClCC(=O)Nc1c(cnn1-c1ccccc1)C#N |
| PP200 | CN(Cc1ccccc1C)C(=O)CCl |
| PP203 | CCN(Cc1ccc(cc1F)C#N)C(=O)CCl |
| PP207 | ClCC(=O)N1c2ccccc2NC(=O)C11CCCC1 |
| PP208 | ClCC(=O)NCc1ccco1 |
| PP209 | Cc1ccc(C)c(NC(=O)CCl)c1 |
| PP214 | ClCC(=O)NCC(=O)N1CCCC1 |
| PP215 | ClCC(=O)Nc1cc(Cl)ccc1-n1cncn1 |
| PP216 | CC(C)n1nccc1NC(=O)CCl |
| PP219 | ClCC(=O)Nc1cc(Cl)ccc1C#N |
| PP221 | CS(=O)(=O)c1ccc(NC(=O)CCl)cc1 |
| PP222 | OC(=O)c1ccc(Cl)cc1NC(=O)CCl |
| PP224 | CC1CCCCN1C(=O)CCl |
| PP225 | CC1Cc2ccccc2N1C(=O)CCl |
| PP232 | CC(=O)Nc1ccc(NC(=O)CCl)cc1 |
| PP48a | COc1ccc(cc1)C(=O)N1CCCN(CC1)C(=O)CCl |
| PP48b | ClCC(=O)N1CCCN(CC1)C(=O)c1ccccn1 |
| PP48c | ClCC(=O)N1CCCN(CC1)C(=O)c1ccc2ccccc2c1 |
| PP48d | ClCC(=O)N1CCCN(Cc2ccc(cc2)C#N)CC1 |
| PP48e | ClCC(=O)N1CCCN(CC1)C(=O)Nc1ccccc1 |
| PP48f | COc1ccc(cc1)S(=O)(=O)N1CCCN(CC1)C(=O)CCl |
| PP48g | COc1ccc(cc1)C(=O)N1CCN(CC1)C(=O)CCl |

**Supplementary Table 2. The diaPASEF method for chemoproteomics data acquisition on a Bruker timsTOF Pro 2.** This diaPASEF method has been formatted to import into the acquisition software timsControl.

| MS Type | Cycle Id | Start IM [1/K0] | End IM [1/K0] | Start Mass [m/z] | End Mass [m/z] | CE [eV] |
|---|---|---|---|---|---|---|
| MS1 | 0 | - | - | - | - | - |
| PASEF | 1 | 1.032 | 1.638 | 800 | 825 | - |
| PASEF | 1 | 0.835 | 1.031 | 600 | 625 | - |
| PASEF | 1 | 0.6 | 0.834 | 400 | 425 | - |
| PASEF | 2 | 1.056 | 1.638 | 825 | 850 | - |
| PASEF | 2 | 0.858 | 1.055 | 625 | 650 | - |
| PASEF | 2 | 0.6 | 0.858 | 425 | 450 | - |
| PASEF | 3 | 1.08 | 1.638 | 850 | 875 | - |
| PASEF | 3 | 0.884 | 1.079 | 650 | 675 | - |
| PASEF | 3 | 0.6 | 0.883 | 450 | 475 | - |
| PASEF | 4 | 1.106 | 1.638 | 875 | 900 | - |
| PASEF | 4 | 0.909 | 1.105 | 675 | 700 | - |
| PASEF | 4 | 0.6 | 0.908 | 475 | 500 | - |
| PASEF | 5 | 1.13 | 1.638 | 900 | 925 | - |
| PASEF | 5 | 0.934 | 1.129 | 700 | 725 | - |
| PASEF | 5 | 0.6 | 0.932 | 500 | 525 | - |
| PASEF | 6 | 1.154 | 1.638 | 925 | 950 | - |
| PASEF | 6 | 0.958 | 1.153 | 725 | 750 | - |
| PASEF | 6 | 0.6 | 0.957 | 525 | 550 | - |
| PASEF | 7 | 1.18 | 1.638 | 950 | 975 | - |
| PASEF | 7 | 0.983 | 1.179 | 750 | 775 | - |
| PASEF | 7 | 0.6 | 0.982 | 550 | 575 | - |
| PASEF | 8 | 1.204 | 1.638 | 975 | 1000 | - |
| PASEF | 8 | 1.006 | 1.202 | 775 | 800 | - |
| PASEF | 8 | 0.6 | 1.005 | 575 | 600 | - |

**Supplementary Table 3. The diaPASEF method for global proteomics data acquisition on a Bruker timsTOF Pro 2.** This diaPASEF method has been formatted to import into the acquisition software timsControl. This method has been optimised by pyDIAD.[10]

| MS Type | Cycle Id | Start IM [1/K0] | End IM [1/K0] | Start Mass [m/z] | End Mass [m/z] | CE [eV] |
|---|---|---|---|---|---|---|
| MS1 | 0 | - | - | - | - | - |
| PASEF | 1 | 0.97 | 1.6 | 687.01 | 710.36 | - |
| PASEF | 1 | 0.82 | 0.97 | 538.27 | 554.96 | - |
| PASEF | 1 | 0.6 | 0.82 | 262.18 | 394.54 | - |
| PASEF | 2 | 0.99 | 1.6 | 710.36 | 736.37 | - |
| PASEF | 2 | 0.85 | 0.99 | 554.96 | 572.28 | - |
| PASEF | 2 | 0.6 | 0.85 | 394.54 | 424.74 | - |
| PASEF | 3 | 1 | 1.6 | 736.37 | 766.07 | - |
| PASEF | 3 | 0.86 | 1 | 572.28 | 589.61 | - |
| PASEF | 3 | 0.6 | 0.86 | 424.74 | 446.91 | - |
| PASEF | 4 | 1.01 | 1.6 | 766.07 | 800.93 | - |
| PASEF | 4 | 0.87 | 1.01 | 589.61 | 606.87 | - |
| PASEF | 4 | 0.6 | 0.87 | 446.91 | 466.76 | - |
| PASEF | 5 | 1.02 | 1.6 | 800.93 | 843.44 | - |
| PASEF | 5 | 0.88 | 1.02 | 606.87 | 625.79 | - |
| PASEF | 5 | 0.6 | 0.88 | 466.76 | 485.77 | - |
| PASEF | 6 | 1.03 | 1.6 | 843.44 | 898.45 | - |
| PASEF | 6 | 0.89 | 1.03 | 625.79 | 645.33 | - |
| PASEF | 6 | 0.6 | 0.89 | 485.77 | 503.79 | - |
| PASEF | 7 | 1.04 | 1.6 | 898.45 | 983.93 | - |
| PASEF | 7 | 0.9 | 1.04 | 645.33 | 665.38 | - |
| PASEF | 7 | 0.6 | 0.9 | 503.79 | 521.48 | - |
| PASEF | 8 | 1.09 | 1.6 | 983.93 | 1398.68 | - |
| PASEF | 8 | 0.93 | 1.09 | 665.38 | 687.01 | - |
| PASEF | 8 | 0.6 | 0.93 | 521.48 | 538.27 | - |
